# ATR16 Syndrome: Mechanisms Linking Monosomy to Phenotype

**DOI:** 10.1101/768895

**Authors:** Christian Babbs, Jill Brown, Sharon W. Horsley, Joanne Slater, Evie Maifoshie, Shiwangini Kumar, Paul Ooijevaar, Marjolein Kriek, Amanda Dixon-McIver, Cornelis L. Harteveld, Joanne Traeger-Synodinos, Douglas Higgs, Veronica Buckle

## Abstract

**Background:** Sporadic deletions removing 100s-1000s kb of DNA, and variable numbers of poorly characterised genes, are often found in patients with a wide range of developmental abnormalities. In such cases, understanding the contribution of the deletion to an individual’s clinical phenotype is challenging.

**Methods:** Here, as an example of this common phenomenon, we analysed 34 patients with simple deletions of ∼177 to ∼2000 kb affecting one allele of the well characterised, gene dense, distal region of chromosome 16 (16p13.3), referred to as ATR-16 syndrome. We characterised precise deletion extent and screened for genetic background effects, telomere position effect and compensatory up regulation of hemizygous genes.

**Results:** We find the risk of developmental and neurological abnormalities arises from much smaller terminal chromosome 16 deletions (∼400 kb) than previously reported. Beyond this, the severity of ATR-16 syndrome increases with deletion size, but there is no evidence that critical regions determine the developmental abnormalities associated with this disorder. Surprisingly, we find no evidence of telomere position effect or compensatory upregulation of hemizygous genes, however, genetic background effects substantially modify phenotypic abnormalities.

**Conclusions:** Using ATR-16 as a general model of disorders caused by sporadic copy number variations, we show the degree to which individuals with contiguous gene syndromes are affected is not simply related to the number of genes deleted but also depends on their genetic background. We also show there is no critical region defining the degree of phenotypic abnormalities in ATR-16 syndrome and this has important implications for genetic counselling.

## Introduction

Cytogenetic, molecular genetic, and more recently, next generation sequencing (NGS) approaches have revealed copy number variations (CNVs) in the human genome ranging from 1 to 1000s kb (Iafrate et al., 2004; MacDonald et al., 2014). CNVs are common in normal individuals and have been identified in ∼35% of the human genome (Iafrate et al., 2004). When present as hemizygous events, in phenotypically “normal” individuals, these imbalances are considered benign; however, CNVs are also amongst the most common causes of human genetic disease and they have been associated with a wide range of developmental disabilities present in up to 14% of the population (Kaminsky et al., 2011).

CNVs have been shown to play an important role in neurodevelopmental disorders including autism spectrum disorder, bipolar disorder, and schizophrenia as well as influencing broader manifestations such as learning disabilities, abnormal physical characteristics and seizures (reviewed in Merikangas et al., 2009). Some CNVs occur recurrently in association with one particular phenotype: for example, deletions within 16p11.2 and/or chromosome 22q are frequently associated with autism, and deletions within 15q13.3 and 1q21.1 are found in schizophrenia. However, the impact of most sporadic CNVs on phenotype is much less clear (Merikangas et al., 2009). Difficulty in interpreting CNVs particularly occurs when they result from complex rearrangements such as those associated with unbalanced translocations, inversions, and imprinting effects.

To understand the principles and mechanisms by which CNVs lead to developmental abnormalities we have simplified the issue by studying the relationship between uncomplicated deletions within the region ∼0.3 to ∼2 Mb in the sub-telomeric region of chromosome 16 and the resulting phenotypes. The 34 individuals studied here (comprising 11 new and 23 previously reported cases) represent a cohort of patients with the α-thalassaemia mental retardation contiguous gene syndrome, involving the chromosomal region 16p13.3, termed ATR-16 syndrome (MIM 141750) (Wilkie et al., 1990).

Individuals studied here have monosomy for various extents of the gene-rich telomeric region at 16p13.3 and all have α-thalassemia resulting from loss of both paralogous α-globin genes. Some patients also have mental retardation (MR), developmental abnormalities and/or speech delay and facial dysmorphism. The most severe cases also manifest abnormalities of the axial skeleton. By precisely defining the endpoints of deletions within 16p13.3 we address whether the associated neurological and developmental defects are simply related to the size of the deletions and the number of genes removed, whether there are critical haploinsufficient genes within this region and whether similar sized deletions always lead to the same phenotypes. Our findings suggest that while the loss of an increased number of genes tends to underlie more severe phenotypic abnormalities, the genetic background in which these deletions occur contributes to the occurrence of mental retardation and developmental abnormalities.

Finally, this subgroup of ATR-16 patients also allowed us to address two long-standing questions associated with large sub-cytogenetic deletions: those of compensatory gene expression and telomere proximity effect (TPE) in cases of telomere repaired chromosomal breakages.

## Methods

### Patients

Ethical approval was obtained under the title “Establishing the molecular basis for atypical forms of thalassaemia” (reference number MREC: 03/8/097) with written consent from patients and/or parents. Here we focus on a cohort of patients with pure monosomy within 16p13.3 to clarify the effect of the deletion. Number of familial cases.

### Fluorescence in situ Hybridisation (FISH)

FISH studies were carried out on fixed chromosome preparations as previously described (Buckle and Rack, 1993) from seven patients (OY, TY, SH, MY, YA using a series of cosmids covering the terminal 2 Mb of chromosome 16p (Supplementary Table 1). FISH studies were also performed using probes specific for the sub-telomeric regions of each chromosome in order to exclude any cryptic chromosomal rearrangements. Subtelomeric rearrangements were detected as previously described (Knight et al., 1997 and Horsley et al., 2001).

### Southern Blotting

Single copy probes labelled with α^32^P-dCTP were synthesized and used to hybridize Southern blots of DNA isolated from transformed lymphoblastoid cells.

### PCR Detection of Chromosomal Deletions

DNA was extracted from mouse/human hybrid cell lines or transformed lymphoblastoid lines. Based on FISH results with chromosome 16 cosmids, primer pairs were designed located at regular intervals across the breakpoint clone. To refine the 16p breakpoint, PCR amplification was performed using normal and abnormal patient hybrid DNA obtained from mouse erythroleukemia (MEL cells) fused to patient cells and selected to contain a single copy of human chromosome 16 generated as previously described (Zeitlin & Weatherall, 1983) as template. A positive PCR indicated the sequence was present; a negative PCR indicated it was deleted. All reactions were carried out in 1.5 mM MgCl_2,_ 200 μM dNTPs, 2U of FastStart Taq DNA polymerase (Roche), 1× FastStart PCR Reaction buffer (Roche) (50mM Tris-HCl, 10 mM KCl, 5 mM (NH_4_)_2_SO_4_, pH 8.3). A list of the primer pairs is shown in Supplementary Table 2.

### Telomere Anchored PCR Amplification

Telomere-anchored PCR was undertaken using a primer containing canonical telomeric repeats in conjunction with a reverse primer specific for the normal 16p sequence (primer sequences provided in Supplementary Table 2). As the telomere repeat primer hybridises at any location in telomere repeats, heterogeneous amplification products were produced. Amplification products were purified and digested with restriction endonucleases BamHI or EcoRI and products ligated into appropriately prepared pBluescript. Resulting colonies were screened for inserts and DNA Sanger sequenced.

### Quantification of gene expression

Total RNA was isolated from Epstein-Barr virus transformed lymphoblastoid cell lines for 10 patients and 20 control individuals using TRI^®^ reagent. cDNA synthesis was performed with the AffinityScript™ kit (Stratagene). Where gene expression was measured by quantitative real-time PCR TaqMan® Gene Expression Assays Applied Biosystems (ABI, www.appliedbiosystems.com) were used. Genes and assay numbers were: *POLR3K* (Hs00363121_m1), *C16orf33* (Hs00430677_m1), *C16orf35* custom assay (Forward; CAACGCCCTCAGCTTTGG, Reverse; GCTGGTGAGGGTCATGTCATC, Probe; CCCCAACCAGCAGC), *LUC7L* (Hs00216077_m1), *AXIN1* (Hs00394718_m1), *MRPL28* (Hs00371771_m1), *TMEM8* (Hs00430491_m1), *NME4* (Hs00359037_m1), *DECR2* (Hs00430406_m1), *Rab11FIP3* (Hs00608512_m1) *ACTB* (Hs99999903_m1). Two custom assays were obtained from Eurogentec (www.uk.eurogentec.com): *MPG* (Forward; 5’-GCATCTATTTCTCAAGCCCAAAG-3’, Reverse; 5’-GGAGTTCTGTGCCATTAGGAAGTC-3’, Probe; 5’-AGTTCTTCGACCAGCCGGCAGTCC-3’) and C16orf9 (Forward; 5’-GGCGGCCCGTTCAAG-3’, Reverse; 5’-GAGCCCACAAGAAGCACA-3’, Probe; 5’-TCCCAGGGAACGCCGGTG-3’). Analysis was performed using the comparative C_T_ Method (ΔΔC_T_) (Livak et al., 2001). Data in Figure 3A were obtained by amplification and Sanger sequencing of informative polymorphisms from genomic DNA and cDNA.

**Figure 1.**
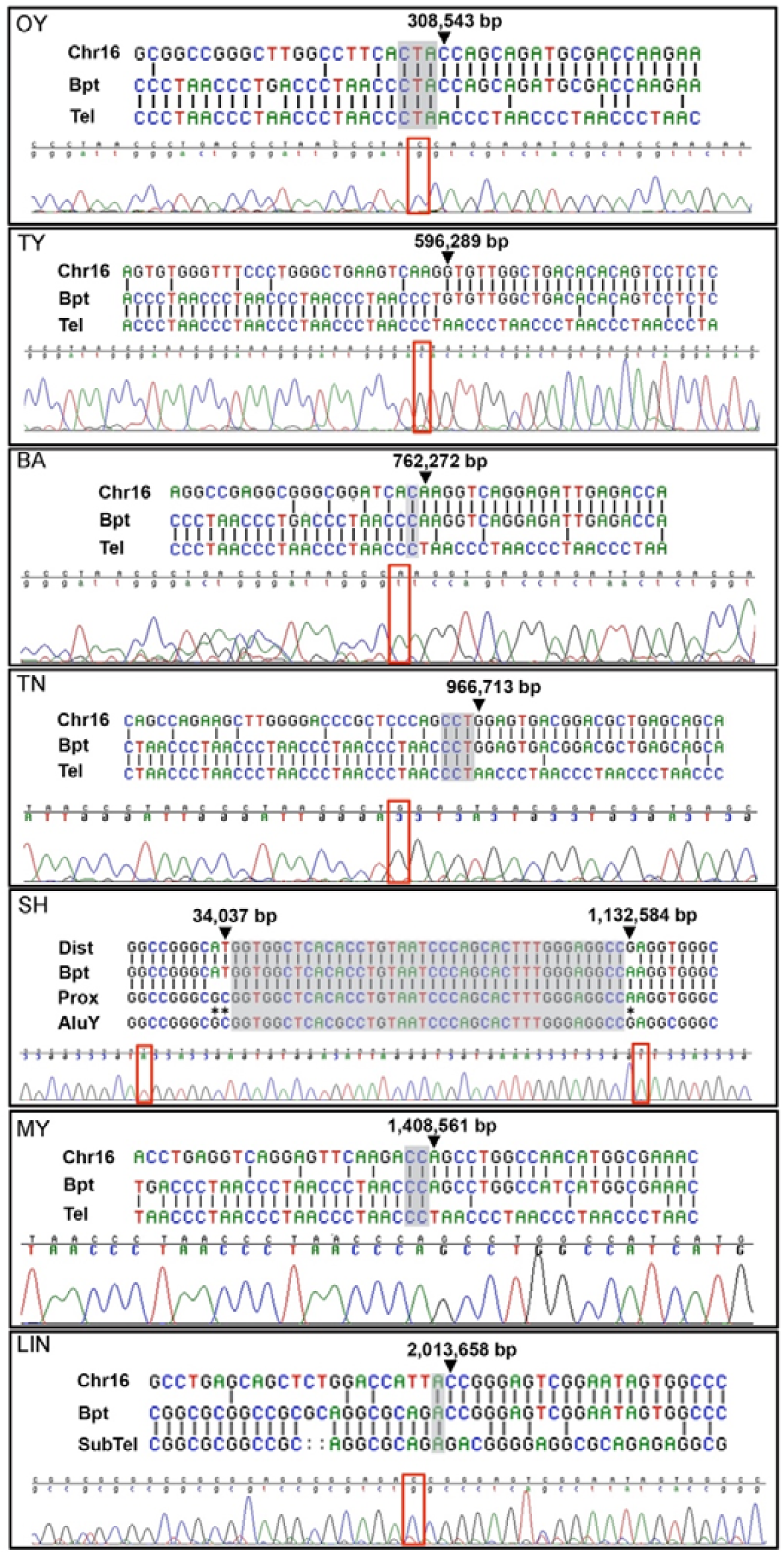
Chromosome 16 breakpoint sequences. DNA sequences at ATR-16 breakpoints. Patient codes are given in the upper left of each panel. For each case alignment of the two normal sequences is shown with sequence from the derivative chromosome (upper) with chromatogram traces traversing each breakpoint (lower). Areas of ambiguity are highlighted with grey boxes and the location of the last unambiguous base pair(s) are denoted by arrowheads and red boxes. Chr16, normal chromosome 16 sequence; Bpt, breakpoint sequence; Tel, telomere repeat sequence; SubTel, subtelomere repeat sequence; Prox, proximal chromosome 16 sequence; Dist, distal chromosome 16 sequence; AluY, AluY repetitive element. Asterisks indicate informative polymorphisms allowing sequence origins to be identified. For patients MY and OY a telomere primer with a mis-matched G nucleotide was used.

**Figure 2.**
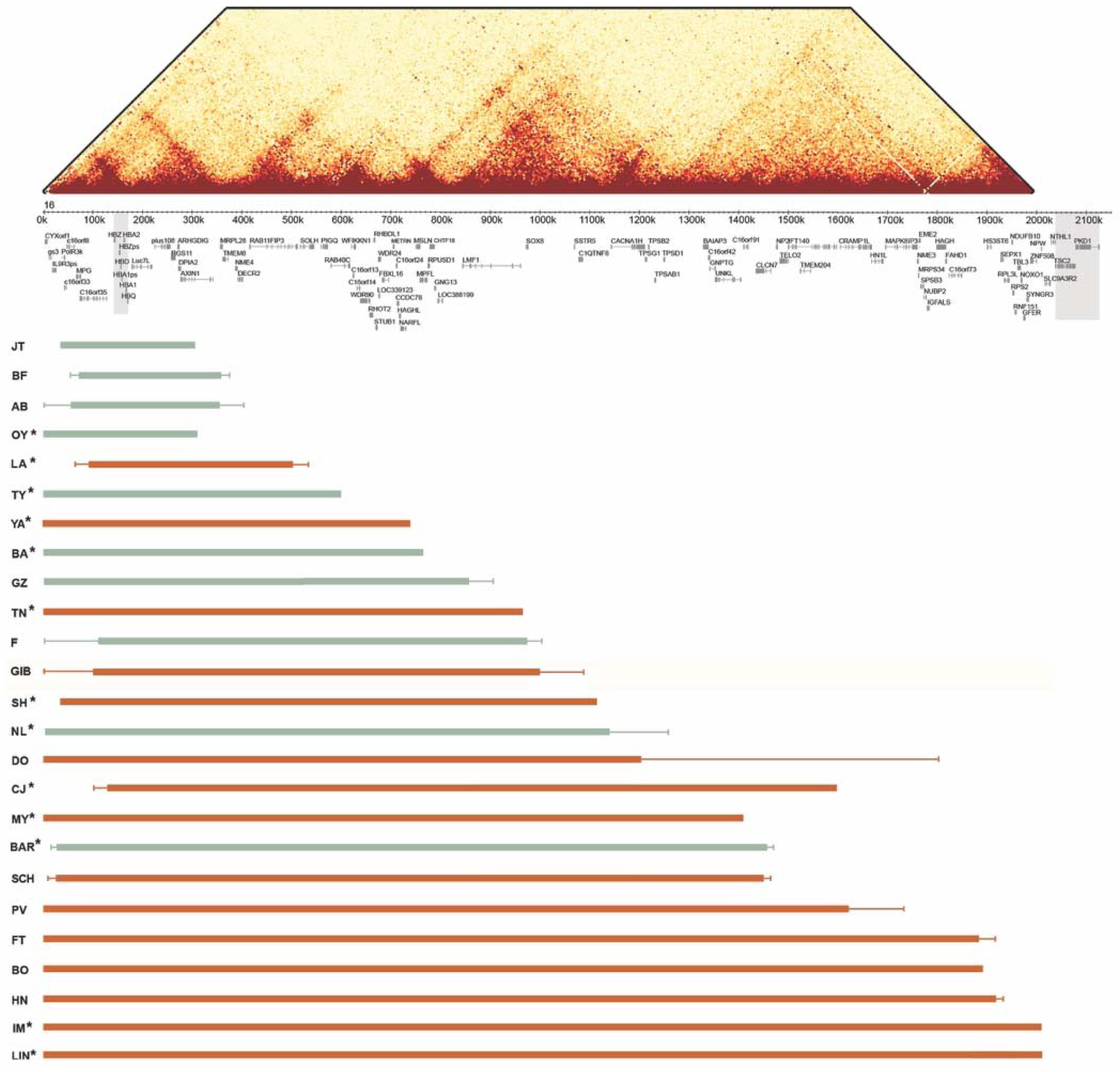
Summary of ATR-16 deletions. Upper: HiC interaction map showing interactions across the terminal 2 Mb of chromosome 16 at a 5 kb resolution in K562 cells (data from Rao et al., 2014). This shows how the ATR-16 deletions detailed in the lower section may impact the genome organisation. Middle: The positions of the alpha-globin cluster and other genes within this region are indicated. The alpha-globin genes and genes that, when mutated, are associated with tuberous sclerosis and adult polycystic kidney disease are shown in shaded boxes. Lower: The extent of each deletion is shown with the patient code (left). Deletions shown in green cause no other abnormalities apart from alpha-thalassemia and those in red cause at least one other abnormality present in ATR-16. Solid bars indicate regions known to be deleted and fine lines show regions of uncertainty. Asterisks indicate individuals whose deletion breakpoints have been cloned or refined in this work.

**Figure 3.**
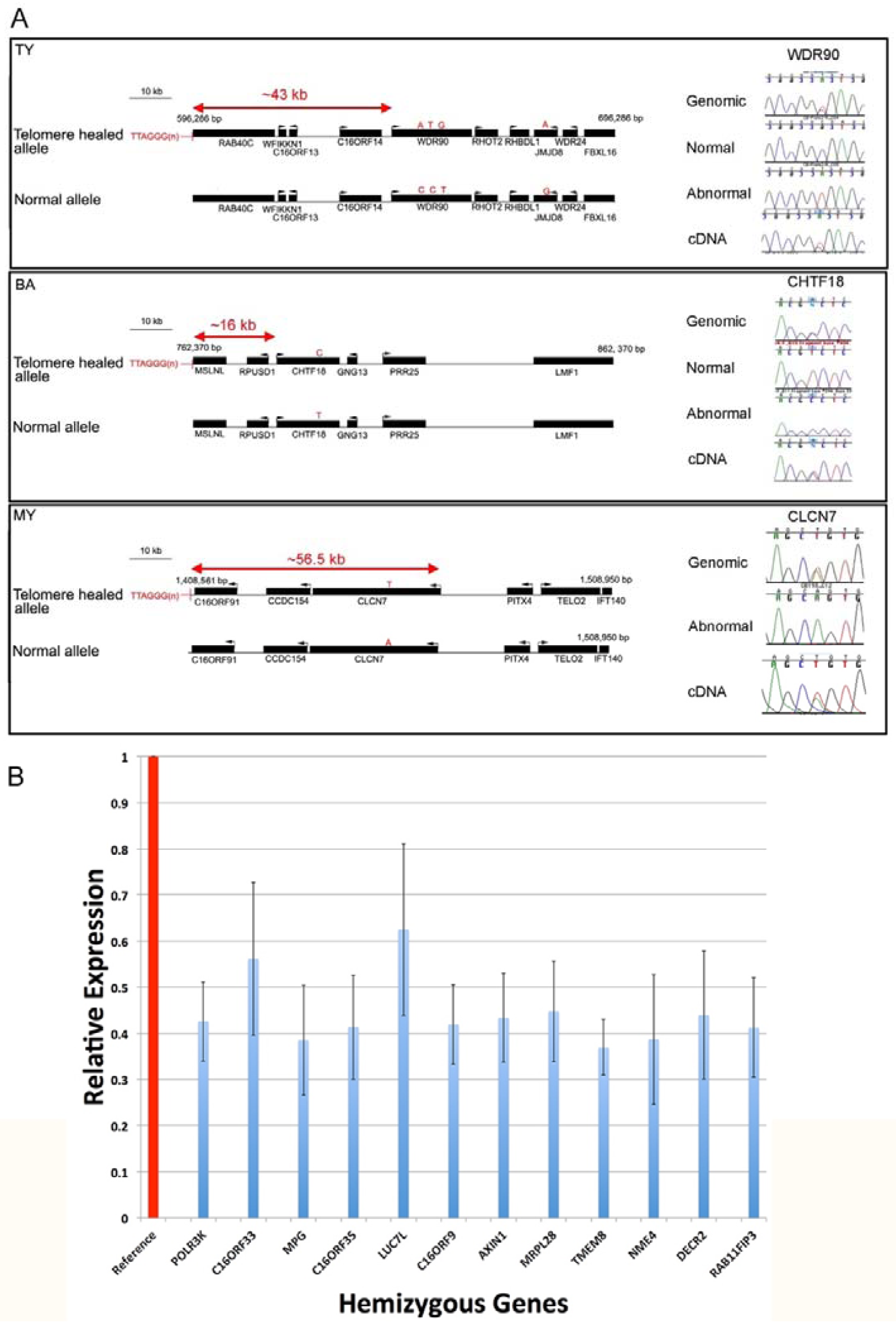
Effect of breakpoints and deletions on gene expression. A. 1 Schematic view of breakpoint positions in 3 patients with nearby expressed polymorphic genes. Genes are represented by black bars and transcription direction is indicated by an arrow. Polymorphic bases are shown by red letters indicating variant alleles and the distance of the promoter of each measured gene from the breakpoint is shown. On the right of each panel chromatograms show the quantity of the allele present in genomic DNA and cDNA from patient lymphoblastoid cells. B. Expression of 12 genes within 500 kb of the tip of the short arm of chromosome 16 in lymphoblastoid cells from 20 normal individuals, shown as reference (red column) and from 10 ATR-16 individuals hemizygous for each gene. Measurements in control cells are normalised to 1 (red column), relative expression in ATR-16 patient cells is shown in blue. Error bars show SD. Gene expression wasmeasured in triplicate and data combined.

### Array Comparative Genomic Hybridisation (aCGH)

DNA from patient LA was tested using an 8 x 60K SurePrint G3 custom CGH + SNP microarray (Agilent) and analysed using Agilent Cytogenomics software 4.0. Genomic DNA from patients NL and BAR was analysed using a custom fine tiling array covering the alpha- and beta-globin gene clusters and surrounding areas was used (Roche NimbleGen, Madison, WI, USA). Array design was based on NCBI Build 36.1 (hg18) and used as previously described (Phylipsen et al., 2012). Genomic DNA from patients SH(Ju) and SH(Pa) were tested using CytoScan HD arrays (Affymetrix) and analysed using Karyoview software. aCGH was performed with genomic DNA from patients CJ, IM and YA using the Sentrix Human CNV370 BeadChip (Illumina) and analysed using GenomeStudio software.

### Whole Genome Sequencing (WGS)

WGS was carried out at Edinburgh Genomics, The University of Edinburgh. The pathogenicity of each variant was given a custom deleterious score based on a six-point scale, (Fu et al., 2013) calculated using output from ANNOVAR (Wang et al., 2010). This was used to prioritise variants present in the hemizygous region of chr16p13.3 in each patient and also genome wide.

## Results

### Clinical Features of ATR-16 Syndrome

All individuals with ATR-16 syndrome have α-thalassemia because at two of the four paralogous α-globin genes are deleted (--/αα) and this manifests as mild hypochromic microcytic anaemia. In combination with a common small deletion involving one α-gene on the non-paralogous allele (--/-α) patients may have a more severe form of α-thlassaemia referred to as HbH disease (Harteveld & Higgs 2010). In addition to α-thalassemia, which is always present, common features of ATR-16 syndrome include speech delay, developmental delay and a variable degree of facial dysmorphism and, in severe cases, abnormalities of the axial skeleton. Newly cloned breakpoint sequences are shown in Figure 1; deletions are shown in Figure 2. Deletions larger than 2000 kb including the *PKD1* and *TSC2* genes lead to severe MR with polycystic kidney disease and tuberous sclerosis respectively (European Polycystic Kidney Disease Consortium, 1994).

Eleven patients from 9 families are reported here for the first time (OY, LA, TY(MI), TY(Mi), YA, SH(P), SH(Ju), NL, CJ, MY and BAR) and we refine the breakpoints in 4 previously reported families (BA, TN, IM, LIN). We define breakpoints at the DNA sequence level in 7 of the 13 families studied (Figure 1), 6 of which have been repaired by the addition of a telomere or subtelomere. In the remaining family (SH) the deletion is interstitial and mediated by repeats termed short interspersed nuclear elements (SINEs).

### Identification of Co-Inherited Deleterious Loci

Three families (LA, YA and TN) have 16p13.3 deletions smaller than 1 Mb and show relatively severe abnormalities. To test whether 16p13.3 deletions of <1 Mb may be unmasking deleterious mutations on the intact chromosome 16 allele in severely affected patients, we performed whole genome sequencing (WGS) where DNA was available (YA and the three affected members of the TN family) and considered only coding variants in the hemizygous region of chromosome 16. However, only common variants (allele frequency >5%) were present (Supplementary Table 3) suggesting the cause(s) of the relatively severe phenotypes in these patients reside elsewhere in the genome. To identity rare variants we considered only those absent from the publicly available databases. This analysis yielded 14 variants shared between the three affected individuals of family TN (Supplementary Table 4). Of these, only one (chr15:64,782,684 G>A) affects a gene likely to be involved in the broader ATR16 phenotypic abnormalities. This change leads to a R12X nonsense mutation in SMAD6, a negative regulator of bone mophogenetic protein (BMP) signalling pathway. Heterozygous mutations in *SMAD6* have been reported to underlie craniosynostosis, speech delay, global developmental delay, fine motor impairment and aortic valve abnormalities with variable penetrance (Timberlake et al., 2016; Luyckx et al., 2019). We checked inheritance of the *SMAD6* R12X variant in members of the TN family where samples were available and found it most likely came from the unaffected grandmother (Individual I,2 in Supplementary Figure 4). A phenotypically normal elder sister also inherited this variant. These findings suggest that coinheritance of this *SMAD6* loss of function variant with the chromosome 16 deletion may lead to the increased severity of the ATR-16 syndrome.

Further evidence the effect of ATR-16 deletions is modified by other loci comes from patients SH(Ju) and SH(Pa) who harbour the same chromosome 16 deletion. Patient SH(Pa) has developmental delay and skeletal abnormalities, however, his mother SH(Ju) does not have craniofacial nor skeletal abnormalities nor developmental delay although she suffers from severe anxiety and depression (see Figures 1,2 and Supplementary Information). Genome wide array comparative genomic hybridisation (aCGH) analysis revealed that both SH(Ju) and SH(Pa) harbour a ∼133 kb deletion on the short arm of chromosome 2 including exons 5 to 13 of *NRXN1* (Supplementary Figure 1). *NRXN1* encodes a cell surface receptor involved in the formation of synaptic contacts and has been implicated in autism spectrum disorder, facial dysmorphism, anxiety and depression, developmental delay and speech delay (e.g. The Autism Genome Project Consortium 2007; Kirov et al., 2008). This finding offers an explanation for the differences in the phenotypic severity of the ATR-16 syndrome affecting these patients as autism differentially affects males and females (see Discussion).

### Chromatin Structure

Recent reports demonstrate chromosomal rearrangements, including deletions, can result in aberrant DNA domain topology and illegitimate enhancer-promoter contact causing gene misexpression (Lupianez et al., 2015). Chromatin contact frequency is shown for the terminal 2 Mb of chromosome 16 in Figure 2A to illustrate the effect of the deletions reported here on the chromatin structure. The deletion in BA removes ∼50% of the self-interacting domain in which *CHTF18, RPUSD1, GNG13*, and *LOC388199* reside thereby potentially removing *cis*-acting regulatory elements of these genes, although the genes themselves remain intact. In the case of CJ, the deletion brings the powerful α-globin enhancer cluster (Hay et al., 2016) into proximity of *CRAMP1L* and may cause its aberrant expression in developing erythroblasts. Although TADs have been reported to be stable structures (e.g. Lupianez et al., 2015), many chromatin contacts are now known to vary in a tissue-specific fashion (Szalaj & Plewczynski, 2018) and therefore it is not possible to predict which genes may be aberrantly expressed in any given tissue as a result of the ATR-16 deletions.

### Compensatory Gene Expression

One explanation for the relatively mild abnormalities in many cases of ATR-16 syndrome with deletions up to 900 kb may be compensatory transcriptional upregulation of the homologues of deleted genes on the undamaged chromosome 16. This has been described as part of the mechanism of genetic compensation, also termed genetic robustness (El-Brolosy and Stainer, 2017). To assay for compensatory gene transcription we used qPCR to measure expression of 12 genes within the terminal 500 kb of chromosome 16 in lymphoblastoid cells from 20 normal individuals and from 10 patients with monosomy for the short-arm of chromosome 16 and found no evidence of compensatory upregulation: transcripts of all deleted genes were present at ∼50% of the normal levels in these cells (Figure 3B). It is possible that other genes in downstream pathways affected by haploinsufficiency may be transcriptionally upregulated, however, the mechanisms underlying this are complex and beyond the scope of this study.

### Telomere Position Effect

Previous work in human cells has shown that telomeres may affect chromatin interactions at distances of up to 10 Mb away from the chromosome ends (Robin et al, 2014) reducing expression of the intervening genes. This phenomenon, termed telomere position effect (TPE) is thought to be mediated by the spreading of telomeric heterochromatin to silence nearby genes. In budding yeast this effect can extend a few kb towards the sub-telomeres, although in some cases yeast telomeres can loop over longer distances (Bystricky et al., 2005) and repress genes up to 20 kb away from the end of the chromosome.

To determine the effect of telomere proximity on genes adjacent to telomere-healed breakpoints we measured their expression relative to the allele present in a normal chromosomal context. To achieve this, we screened them for informative single nucleotide polymorphisms (SNPs) in EBV transformed lymphoblastoid cells generated from ATR-16 patients. The phase of polymorphisms was established using mouse erythroleukemia (MEL) cells fused to patient cells and selected to contain a single copy of human chromosome 16, generated as previously described (Zeitlin & Weatherall 1983). Expressed coding polymorphisms were present in genes whose promoters are <60 kb away from breakpoints in 3 patients: TY, MY and BA.

For TY the nearest gene expressed in lymphoblastoid cells containing a coding polymorphism is *WDR90*, the promoter of which is ∼43.1 kb from the abnormally appended telomere (Figure 3A). For BA, *CHTF18* is the closest expressed polymorphic gene with the promoter ∼16.3 kb away from the breakpoint. For MY, *CLCN7* is the closest gene expressed in lymphoblastoid cells to contain a polymorphism, the promoter of this gene is ∼56.1 kb away from the telomere stabilised lesion. To determine whether either allele of each of these 3 genes is silenced we prepared genomic DNA and cDNA from each cell sample and Sanger sequenced amplified fragments containing informative polymorphisms. We compared peak heights of polymorphic bases in chromatograms derived from cDNA and genomic DNA. None of the alleles assayed in the three patients tested showed any evidence of a repressive effect (Figure 3A).

## Discussion

We characterised deletions leading to simple monosomy of the short arm of chromosome 16 that cause ATR-16 syndrome. Many ATR-16 syndrome patients suffer from neurodevelopmental abnormalities and one of the main questions in this disease, and in the study of CNVs in general, is how deletion size relates to phenotypic abnormalities. The monosomies analysed here show the likelihood and severity of neurological and developmental abnormalities increases with deletion size, however, there is no clear correlation.

The deletions in patients reported here range from ∼0.177 Mb to ∼2 Mb. Previous studies suggest the critical region leading to abnormalities in addition to α-thalassemia is an 800 kb region between ∼0.9 and ∼1.7 Mb from the telomere of chromosome 16p (Harteveld et al., 2007) and *SOX8* has been proposed as the critical haploinsufficient gene (Pfeifer et al., 2000). However, a report of a family with no developmental delay nor MR harbouring a 0.976 Mb deletion, suggests deletions *SOX8* may not lead to MR with complete penetrance and any “critical region” for MR starts after this point (Bezerra et al., 2008; Family “F” in Figure 2). Supporting this we report patients NL and BAR, who have deletions of ∼1.14 Mb and ∼1.44 Mb respectively and show no abnormalities beyond α-thalassemia.

By contrast, we find LA (deletion ∼408 kb) has speech delay and YA (deletion ∼748 kb), has speech and developmental delay and facial dysmorphism (Figure 2). Family members of YA also have omphalocele, umbilical hernia and pyloric stenosis suggesting there are other loci rendering YA susceptible to developmental abnormalities. BA (deletion ∼762 kb), who has a similarly sized deletion to YA, has developmental delay but no other abnormalities. Four other patients with deletions <1 Mb (YA, TN(Pa), TN(Pe) and TN(Al)) have speech delay and facial dysmorphism. This suggests the risk of developmental and neurological abnormalities arises from much smaller terminal chromosome 16 deletions (∼400 kb) than previously reported.

In SH(Pa) we have identified a strong candidate for the discordant abnormalities: SH(Pa) also has a deletion of *NRXN1*, disruptions of which cause autism and a range of neurological disorders. There is a higher incidence of autism in males than in females, with a ratio of 3.5 or 4.0 to 1 (reviewed in Volkmar et al., 2004). This phenomenon is also specifically found in individuals with autism resulting from rearrangements of *NRXN1*: Kirov and colleagues (2008) reported two affected siblings who inherited a deletion of *NRXN1* from their unaffected mother. It is therefore possible Sh(Ju) is protected by her gender from the effects of *NRXN1* disruption while the neurological and skeletal abnormalities in Sh(Pa) arise from the complex interaction of *NRXN1* perturbation with his gender and coinheritance of the 16p13.3 deletion. Abnormalities in siblings of YA and BF (Heireman et al., 2019) also suggest there may be other predisposing genes. Such loci compromise genetic robustness proposed to minimise the effect of deletions and loss of function mutations (El-Brolosy and Stainer, 2017). Another example of this is the SMAD6 R12X nonsense mutation present in all three affected members of family TN. Some patients with loss of function mutations in *SMAD6* have neurological abnormalities (Timberlake et al., 2016) while others have not (Luyckx et al., 2019), suggesting variable penetrance. Our analysis shows there are no likely pathogenic variants on the hemizygous region of chromosome 16 in TN, suggesting modifying loci are present elsewhere in the genome. These may be rare variants (such as those identified in the TN and SH families) or common variation; a recent study that shows that common genetic variants (allele frequency >5% in the general population) contribute 7.7% of the variance of risk to neurodevelopmental disorders (Niemi et al., 2018), highlighting the complexity of this area.

Together these observations suggest that monosomy for 16p13.3 unmasks the effects of other variants genome-wide. This is supported by findings in SCH who has a very similar deletion to BAR and may be more severely affected owing to the presence of other CNVs (Scheps et al., 2016). At the other end of the spectrum, large ATR-16 deletions may be associated with relatively mild abnormalities. In LIN (16p13.3 deletion ∼2000 kb) there are no abnormalities of the axial skeleton and very mild facial dysmorphism. Similarly, in the case of IM (deletion size ∼2000 kb), facial abnormalities are very mild and there is no evidence of language delay. Here we propose chromosome 16p13.3 deletions larger than 400 kb predispose to MR and associated developmental abnormalities, however, we find no evidence for critical regions that incrementally worsen ATR-16 syndrome abnormalities.

We could not detect compensatory up-regulation of the homologues of deleted genes. Recently, a case of ATR-16 was reported with a ∼948 kb deletion and who presented with a neuroblastoma *in utero* (Quadrifoglio et al., 2016). These authors speculate that haploinsufficiency of the tumour suppressor *AXIN1* may have contributed to the neuroblastoma. Our finding that the remaining *AXIN1* allele shows no compensatory expression supports this hypothesis.

Terminal chromosome deletions are the most common subtelomeric abnormalities (Ballif et al., 2007). The 16p deletions reported here are among the most common terminal deletions along with 1p36 deletion syndrome, 4p terminal deletion (causing Wolf-Hirschhorn syndrome), 5p terminal deletions (causing Cri-du-chat syndrome), 9q34 deletion syndrome and 22q terminal deletion syndrome. Despite their impact on human health the mechanisms and timing underlying telomeric breakage remain unknown. Findings of terminal deletions of 16p reported and reviewed here and smaller deletions previously reported by our laboratory (Flint et al., 1994) compared to more complex rearrangements at 1p, 22q and 9qter implies different chromosomes are predisposed to different breakage and rescue mechanisms. ATR-16 deletions are equally likely to have arisen on the maternal or paternal chromosome. There is no evidence that the parental origin affects the phenotypic severity of the ATR-16 syndrome, suggesting imprinting does not play a role in ATR-16 pathogenesis.

The presence of high- and low-copy-number repeats at breakpoints may play a role in stimulating the formation of non-recurrent breakpoints (Gu et al., 2008). Low copy repeats (LCRs) are also mediators of nonallelic homologous recombination (Stankiewicz and Lupski, 2002) and could be involved in chromosome instability leading to terminal deletion. Following breakage, chromosomes can acquire a telomere by capture or *de novo* telomere addition, which is thought to be mediated by telomerase and this is stimulated by the presence of a telomeric repeat sequence to which the RNA subunit of telomerase can bind (reviewed in Hannes et al., 2010). We found 5 out of 6 telomere healed events share microhomology with appended telomeric sequence. This is the same ratio (5 out of 6 breakpoints with microhomology) described by Flint and colleagues and supported by Lamb and colleagues (1 out of 1) giving a total 12 out of 13 reported telomere-healed breaks characterised on 16p13.3 share microhomology with appended telomere sequences, strongly suggesting a role for internal telomerase binding sites (Yang et al., 2011). It may also be that telomerase binding to internal binding sites may inappropriately add telomeres and thereby contribute to the generation of the breakpoints.

The lack of evidence for TPE in silencing gene expression is surprising and at variance with previous findings (Stadler et al., 2013), which show that TPE can influence gene expression at least 80 kb from the start of telomeric repeats. However, TPE is likely to be context and cell type dependent. Additionally, because of the lack of informative expressed polymorphisms in the patients studied here it was not possible for us to assay expression of genes immediately adjacent to telomeres and a more comprehensive screen may reveal TPE mediated gene silencing closer to the telomere. Additionally, when the area of chromatin interaction (visualised by HiC) is considered (Figure 2), contact domains for many genes adjacent to chromosomal breaks are severely disrupted. This is likely to include the loss of *cis*-acting regulatory elements and may bring the genes under the control of illegitimate regulatory elements (Franke et al., 2016). Therefore, it is likely that genes adjacent to breakpoints would be incorrectly spatiotemporally expressed.

This work substantially increases the number of fully characterised cases of ATR-16 syndrome reported and provides a uniquely well characterised model for understanding how sporadic deletions giving rise to extended regions of monosomy may affect phenotype. The findings show larger deletions have a greater impact but importantly our analysis suggests there is no critical region defining the degree of phenotypic abnormalities this has important implications for genetic counselling. Analysis of patients with uncomplicated deletions also revealed unexpected background genetic effects that alter phenotypic severity of CNVs.

## Acknowledgements

The authors thank Markissia Karagiorga-Lagana, MD, ex Director, Thalassaemia Unit, “Aghia Sophia” Childrens Hosital Athens Greece for referring case BAR. This work was supported by the Medical Research Council grant number: (MC_uu_12009).

## Authorship Statement

DRH and VB conceived the project. CB, VB and DRH wrote the paper. CB, JS, CLH, JB, EM, SWH, VB, SK and AD-M, performed experimental work. JTS, PO, MK, AD-M assessed patients and clinical data. All authors commented on the manuscript.

