## Supplementary Information for "ATR16 Syndrome: Mechanisms Linking Monosomy to Phenotype"

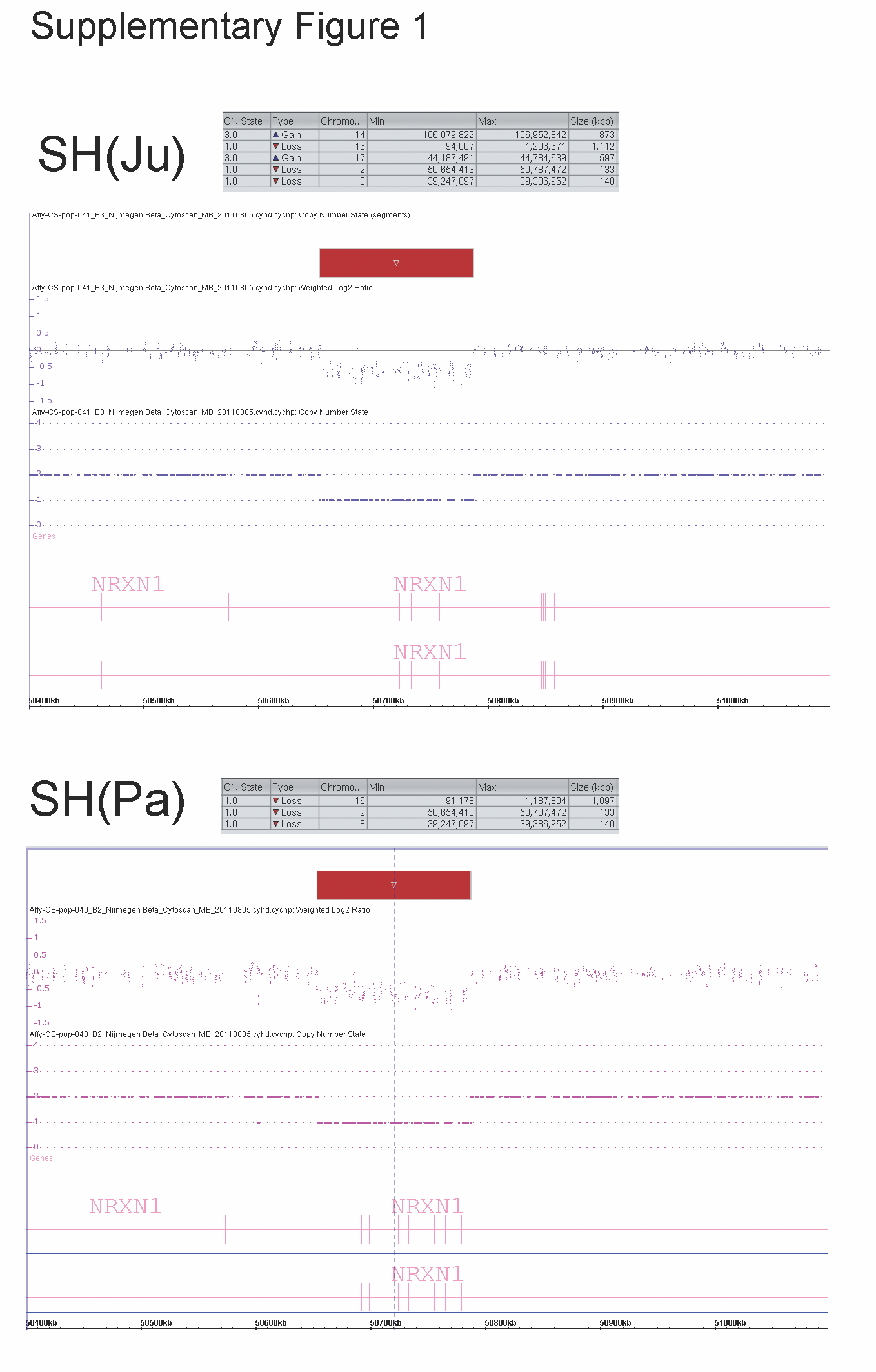

**Supplementary Figure 1:** Copy number variations (CNVs) identified in SH(Ju) and SH(Pa). Tables show CNVs called by Affymetrix Cytoscan software and figures below show the log2 ratio of probes and the copy number state for the chromosome 2 deletion encompassing exons 5 to 13 of *NRXN1*. Both individuals also harbour deletions of chromosomes 8 and 16. The chromosome 8 deletion disrupts the pseudogene *ADAM5* and *tMDC* encoding uncharacterised protein ENSP00000328747. The chromosome 16 deletion underlies the ATR-16 syndrome in these patients.

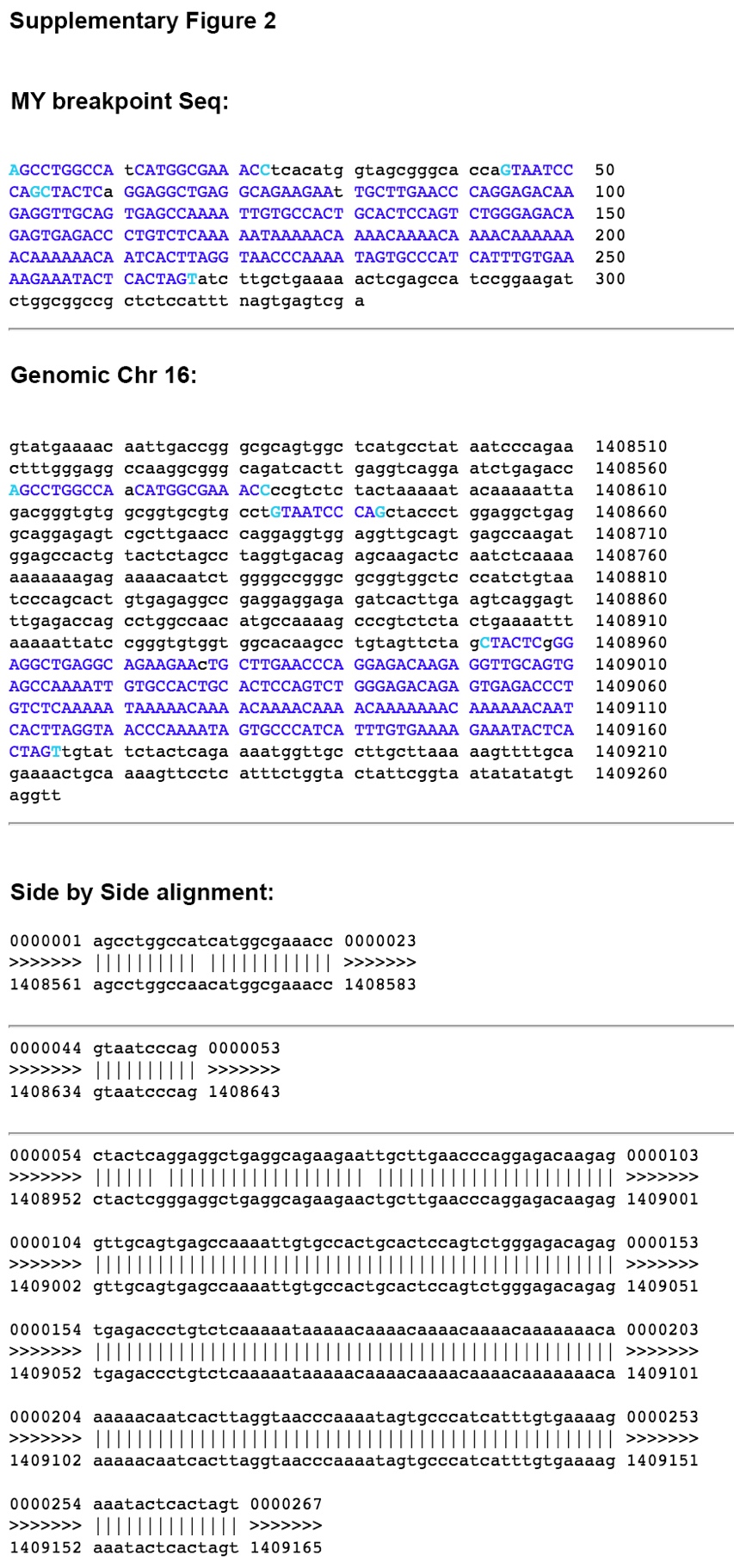

**Supplementary Figure 2:** DNA sequence of breakpoint in patient MY (upper) shown in comparison to genomic sequence with MY breakpoint sequence highlighted in blue (middle) and the side by side alignment of these sequences (lower). The breakpoint present in MY is complex and there is rearrangement of the Alu repeat associated with breakpoint.

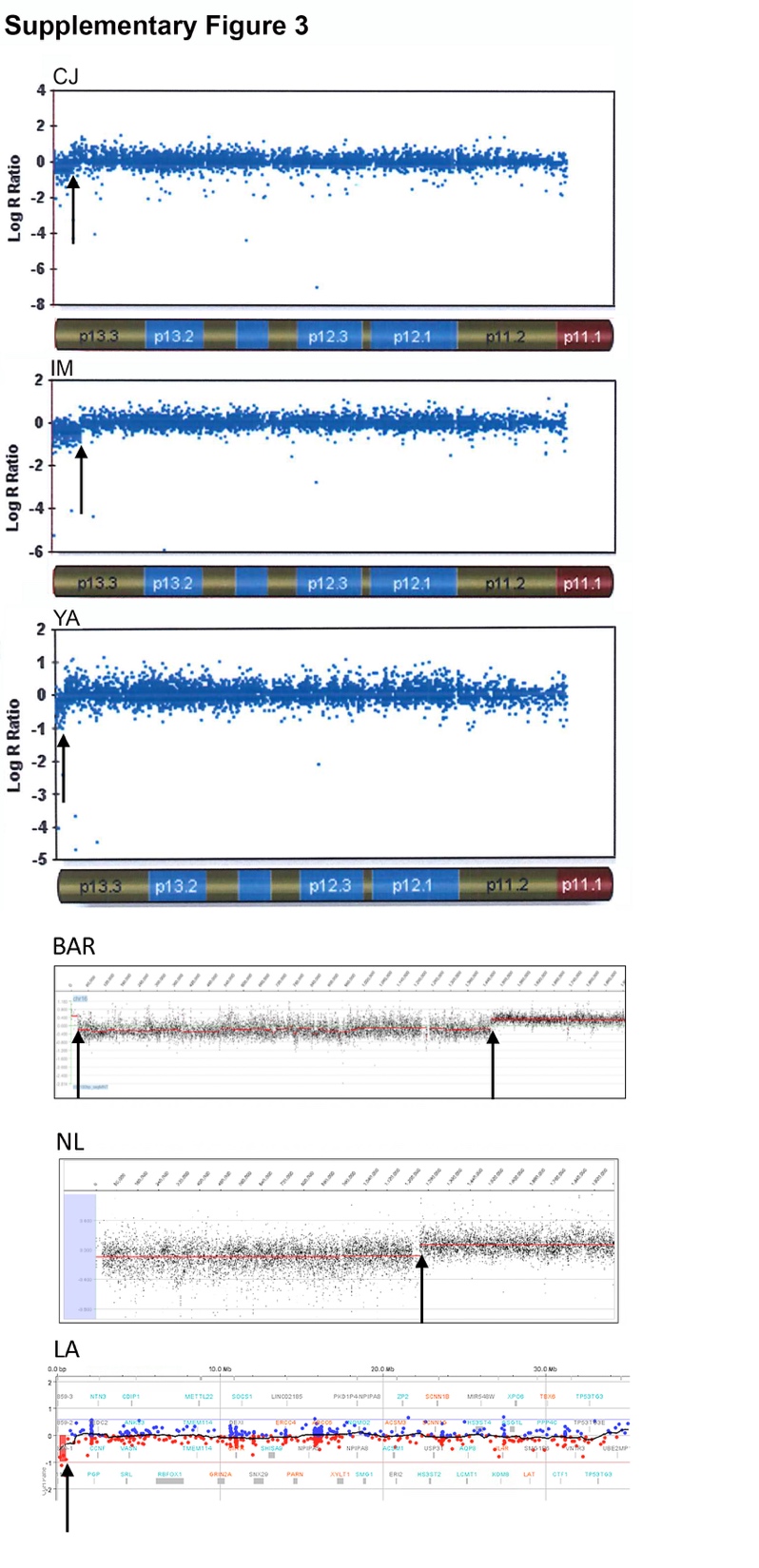

**Supplementary Figure 3:** Array CGH data from patients CJ, IM, YA, BAR, NL and LA showing the LogR ratios of individual probes, arrows indicate loci at which analysis software identified the breakpoints.

**Supplementary Figure 4**

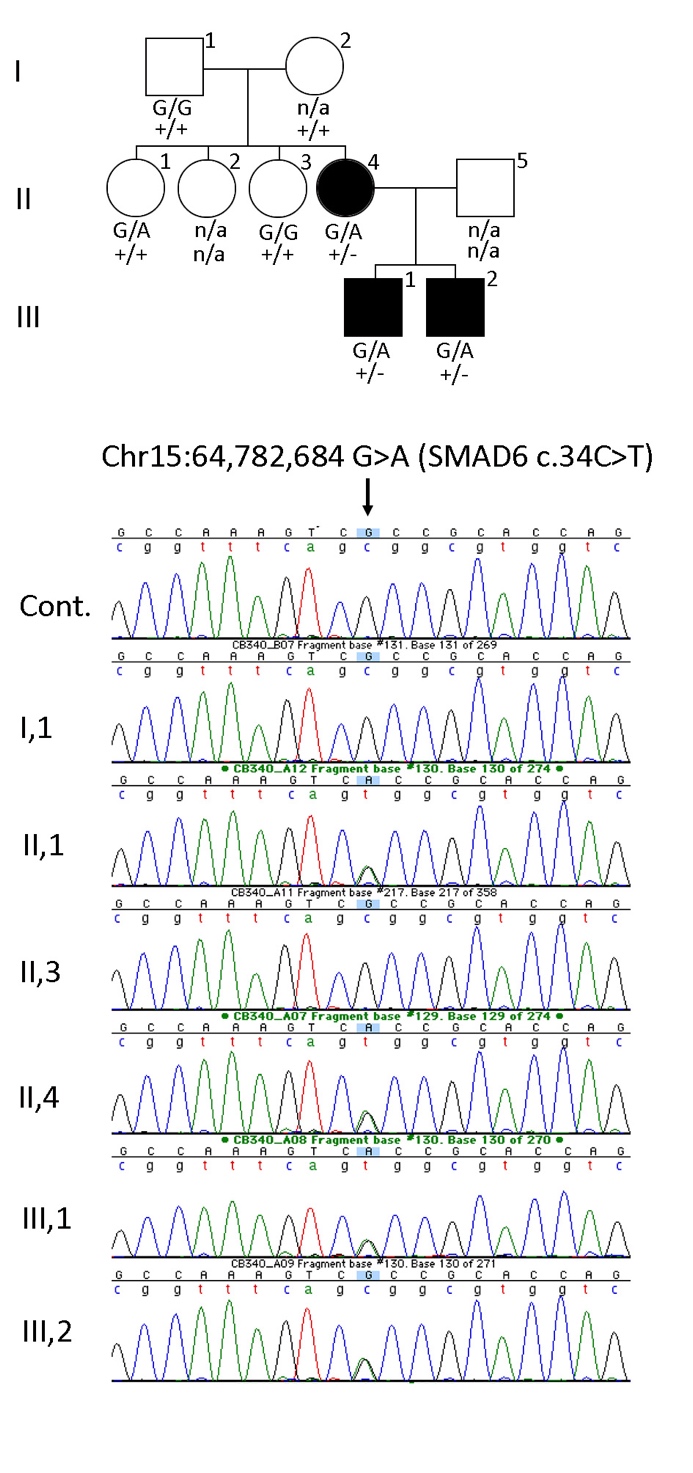

**Supplementary Figure 4.** Identification of SMAD6 change in Family TN. Affected family members are shown with filled symbols and unaffected individuals are represented with open symbols. Below each symbol the genotype at chr15:64,782,684 is written on the upper line, G is reference and A is the variant leading to the R12X nonsense change in *SMAD6*. The lower line of text indicates the ATR16 deletion in this family, + indicates an intact chromosome 16 and – indicates the deleted allele. Because two siblings carry the *SMAD6* variant this is likely to have been inherited from the grandmother (I,2). Coinheritance of these variants may have led to the relatively severe ATR16 abnormalities present in this family. The lower part of the figure shows chromatograms for a control individual (Cont.) and family members as indicated.

**Supplementary Table 1** FISH Probes

| **Probe** | **Coordinates** | **OY** | **TY** | **SH** | **MY** | **LIN** | **YA** | **CJ** |
| --- | --- | --- | --- | --- | --- | --- | --- | --- |
| Telomere Assay | na | N | nd | nd | N | nd | N | N |
| **16p cosmid** |  |  |  |  |  |  |  |  |
| CRA36 | 52706-99963 bp | - |  |  |  |  |  | + |
| HSGG4 | 97541-111766 bp |  | - |  | - |  | - |  |
| HSGG1 | 129137-172194 bp | - | - |  | - |  | - | - |
| Cos12 | 158777-203194 bp |  | - |  |  |  |  | - |
| C314G4 | 243192-285276 bp | - |  |  |  |  |  |  |
| C419C1 | 265315-316860 bp | +/- | - |  |  |  |  | - |
| C338B10 | 297847-340834 bp | + |  |  |  |  |  |  |
| C415C1 | 332201-381990 bp | + |  |  | - |  |  |  |
| C356B8 | 485020-530746 bp | + |  |  |  |  |  |  |
| C366D1 | 521427-565641 bp |  | - |  |  |  |  |  |
| C407A10 | 557634-598294 bp |  | - |  |  |  |  |  |
| C338H10 | 580029-610790 bp |  | +/- |  | - |  |  |  |
| C398G5 | 594663-640361 bp | + | + |  |  |  |  |  |
| C444G9 | 708336-751169 bp |  |  |  |  |  | - |  |
| C335H7 | 737067-783265 bp |  |  |  |  |  | + |  |
| C360B4 | 851308-895707 bp |  |  |  | - |  |  |  |
| C313F9 | 1025418-1072203 bp | + |  |  |  |  |  |  |
| C349E11 | 1068736-1111710 bp |  |  | - | - |  |  | - |
| C344F5 | 1122769-1160074 bp |  |  | + |  |  |  |  |
| C320E12 | 1172104-1208298 bp |  |  |  |  |  |  | - |
| C399E4 | 1359308-1404844 bp |  |  |  | - |  |  | + |
| C312E8 | 1395863-1429931 bp |  |  |  | + |  |  |  |
| C305C8 | 1461998-1505438 bp | + |  |  | + |  |  | + |
| C371H6 | 1755695-1799062 bp |  |  |  | + |  |  |  |
| C439A6 | 1882600-1923854 bp | + |  |  |  |  |  |  |
| 910O19 | 1965888-2004708 bp |  |  |  |  | +/- |  |  |
| 1308N18 | 1996694-2036573 bp |  |  |  |  | + |  |  |
| 2171017 | 2038309-2078584 bp |  |  |  |  | + |  |  |

Presence of signals on both chromosome 16 homologues is indicated by “+”, “-“ indicates signal absent on one chromosome 16 homologue, “+/-“ indicates weaker signal on one homologue compared to the other. The top row indicates patients who were analysed for subtelomeric rearrangements using a telomere assay (Knight et al., 1997 and Horsley et al., 2001): N, no rearrangement; nd, not done; na, not applicable.

**Supplementary Table 2** Oligonucleotides used to clone breakpoints

| Patient | Forward primer | Reverse primer |
| --- | --- | --- |
| SH | 5'-AGATACATGCTCCCAGTCTCA-3' | 5'-CGTATATCTGGTCTCTATCTTC-3' |
| OY | 5'-CAAAGCACGCATCCATAGGC-3' | 5'-CCCTAACCCTGACCCTAACCC-3' |
| LN | 5'- GCAGAGGGAGAGCAGGTCTCAG-3' | 5'-CCCTAACCCTGACCCTAACCC-3' |
| MY | 5'- CTGAAGGACTTGGCTGGTGGAT-3' | 5'-CCTAACCCTGACCCTAACCC-3' |
| BA | 5'-GAGCAAAGTACACAAACTGGGTGAC-3' | 5'-CCTAACCCTGACCCTAACCC-3' |
| TN | 5'-AACTGGCCTTGTCTGTGCCTTAAGCT-3' | 5'-CCCTAACCCTGACCCTAACCC-3' |
| TY | 5'-CCTACCACCAGCAAGAACGGA-3' | 5'-CCTAACCCTGACCCTAACCC-3' |

**Supplementary Table 3** Deleterious Chromosome 16 Variants

| Patients* | Gene^#^ | Variant^%^ | Identifier^ | Freq^$^ | ANNOVAR^@^ |
| --- | --- | --- | --- | --- | --- |
| TN(Al, Pa, Pe) | MRPL28 | H27Y | rs3194151 | 11% | 3/6 |
| TN (Al,Pe) | PICQ | G523S | rs7187227 | 14% | 3/6 |
| TN(Al) YA | PICQ | T14A | rs2071979 | ~49% | 2/6 |
| TN(Al, Pa,Pe) YA | POLR3K | S24A | rs3194151 | 100% | 2/6 |
| TN(Pa) | PDIA | T286M | rs2685127 | 8% | 3/6 |
| TN(Pa) | CHTF | S63F | rs2277902 | 5% | 1/6 |
| TN(Pa) | PRR25 | T92S | rs1005190 | 41% | 1/6 |
| YA | NPRL3 | R158fs | rs35963490 | 33% | 6/6 |
| YA | RGS11 | G499A | rs9806942 | 15% | 3/6 |
| YA | RHOT | R425C | rs3177338 | 33% | 3/6 |
| YA | RHOT | R245Q | rs1139897 | 33% | 2/6 |
| YA | RGS11 | T728C | rs739999 | 31% | 1/6 |
| YA | WDR90 | H899Q | rs45613635 | 34% | 1/6 |

*Codes for patients harbouring each of the changes listed are shown in this column. ^#^Gene symbols are used. ^%^ Effects of variants on coding sequence.^$^ Allele frequencies as a population average, data from dbSNP. ^@^ The pathogenicity of each variant was given a custom deleterious score based on a six-point scale, (Fu et al., 2013) calculated using output from ANNOVAR (Wang et al., 2010).

**Supplementary Table 4**. Shared novel variants in 3 affected members of family TN

| **Chr^#^** | **Position (bp)** | **Gene** | **Variant^$^** | **ANNOVAR^%^** | **Associated abnormalities^** |
| --- | --- | --- | --- | --- | --- |
| 1 | 1336598 | PRAMEF27 | Q144R | 1/6 | na |
| 1 | 26671545 | CRYBG2 | L524fs | 6/6 | na |
| 2 | 71190373 | ATP6V1B1 | R331W | 4/6 | Renal tubular acidosis with progressive nerve deafness. |
| 3 | 141163642 | ZBTB38 | K804N | 5/6 | Potential role in human height variation (nearby GWAS association). |
| 4 | 17061677 | CLCN3 | A290S | 4/6 | Loss of hippocampus in mouse KO. |
| 5 | 179068881 | C5orf60 | splicing | 6/6 | na |
| 8 | 125052168 | FER1L6 | G837D | 3/6 | na |
| 11 | 1018248 | MUC6 | T1518I | 2/6 | Gastric cancer |
| 11 | 62293804 | AHNAK | F2695L | 2/6 | Neuroblastoma |
| 13 | 103718412 | SLC10A2 | K63fs | 6/6 | Chrohn’s disease, bile malabsorbtion. |
| 14 | 33014538 | AKAP6 | L227V | 5/6 | na |
| 15 | 66995630 | SMAD6 | R12X | 6/6 | Aortic Valve Disease, Craniosynostosis, developmental delay. |
| 19 | 59010556 | SLC27A5 | V483M | 6/6 | na |
| 20 | 5170758 | CDS2 | P406A | 4/6 | na |

^#^Chromosome. ^$^The effect of each variant on the protein is shown. ^%^The custom pathogenicity score attributed to each variant (see Supp Table 3 notes). ^Annotation present in the OMIM database associated with each gene, na indicates no information in OMIM. The pathogenicity of each variant was given a custom deleterious score based on a six-point scale, (Fu et al., 2013) calculated using output from ANNOVAR (Wang et al., 2010).
